# AKNA drives neural stem cell fate transition through differential localization and coordinating the modulation of chromatin from H3K27me3 to H3K27ac

**DOI:** 10.1101/2025.09.15.676257

**Authors:** Soumen Manna, R Kirtana, Tirthankar Baral, Jagdish Mishra, Piyasa Nandi, Subhajit Chakraborty, Niharika Mahto, Ankan Roy, Prahallad Mishra, Bhagyashree Pradhan, Pujarini Dash, Samir Kumar Patra

**Affiliations:** Epigenetics and Cancer Research Laboratory, Biochemistry and Molecular Biology Group, Department of Life Science, National Institute of Technology, Rourkela, Odisha, 769008, India And ICMR Collaborating Centre of Excellence (ICMR-CCoE), Department of Life Science, National Institute of Technology, Rourkela, Odisha, 769008, India

**Keywords:** AT-hook, AKNA, epigenetics, neuroblastoma, Stemness, differentiation, chromatin dynamics

## Abstract

Neuronal stem cells (NSCs) play pivotal role in adult neurogenesis, however, detail mechanisms of obtaining pluripotency or undergoing differentiation remain unknown. Herein, how AT-hook protein AKNA regulates pluripotency and stemness in neuroblastoma cells is demonstrated by gene knockdown, immunofluorescence, chromatin immunoprecipitation (ChIP), localization of AKNA, signaling interactions and transcriptional activity. AKNA was abundant and mostly nuclear during induction of pluripotency and its knockdown reduced stemness, even in presence of other pluripotency factors, including OCT4 and SOX2. During the course of induction of differentiation, AKNA remain localized in the cytosol, essential for regulated differentiation. Cytosolic retention of AKNA is possibly driven by FAK signaling. In the nucleus, AKNA promotes KDM6B demethylase for H3K27me3 demethylation to H3K27, and subsequently promotes CBP/p300 to acetylate H3K27 to H3K27ac deposition on promoter region of respective genes to trigger their transcription. Data obtained from knockdown and overexpression of AKNA and KDM6B further reinforce its importance that, they physically interact to drive pluripotency and stemness. These findings establish AKNA as a critical regulator for NSCs fate determination in association with epigenetic modifiers and signaling pathway, offering potential targets for neuroblastoma therapies and regenerative medicines for neurodegenerative diseases.

## Introduction

AT-hook containing proteins are a class of protein characterized by small AT-motif, which is sufficient to bind to the DNA minor groove and plays important roles in regulating chromatin remodeling and gene expression from plants to animals (Yun et al. 2012)(Aravind and Landsman 1998). One of such protein is AKNA that garnered attention due to its roles in immune system regulation and development. Earlier it was characterized as nuclear localized transcription factor (TF) in CD40 and CD40L promoters during secondary immune response (Huth et al. 1997)(Reeves and Nissen 1990)(Siddiqa et al. 2001). Later, AKNA was described as a centrosomal protein which regulates neurogenesis by modulation of microtubule organizations. It also regulates the epithelial to mesenchymal transition in neural stem cells delamination in subventricular zone of brain (Camargo Ortega et al. 2019). Shifting of transcription regulatory proteins from nucleus to cytosol or vice versa is an established phenomenon for multifaceted functions. For example, Inhibitor of differentiation 1 (ID1), helix-loop-helix protein is a stemness like gene, helps in survival and proliferation. ID1 is re-localized in cytoplasm after treatment of retinoic acid in SH-SY5Y cells (Cheung et al. 2009). Coordination between chromatin modifiers, TFs and DNA architecture modifying proteins regulates gene expression. Among the chromatin modifiers histone methyl transferases (HMTs) and demethylases (KDMs) are some of the important enzymes has taken the leads with their crosstalk with different TFs (Manna et al. 2023b).

Remarkable capacity of TFs to cell fate determination depends on chromatin remodeling at target loci by recruiting specific epigenetic modifiers. Many AT-hook containing proteins themselves acts as a chromatin remodelers like ALL1, SWI/SNF, TAFII250, Ash1 etc. (Alver et al. 2017)(Collins et al. 2019)(Jacobson et al. 2000)(Dorafshan et al. 2019)(Aravind and Landsman 1998). During pluripotent stem cells (PSCs) self-renewal and differentiation, epigenetic regulation preserves cell identity by shaping unique epigenetic landscapes. Bivalent domains, marked by both the repressive histone H3K27 trimethylation (H3K27me3) and the activating H3K4 trimethylation (H3K4me3), are uniquely positioned at lineage-specific genes in embryonic stem cells (ESCs), poising them for precise activation or repression during fate transitions. Research indicates that deposition of H3K4me3 tends to be often precede demethylation of H3K27me3, pre-activating genes but repressing them until H3K27me3 is removed by KDM6 family demethylases (e.g., KDM6A/UTX, KDM6B/JMJD3) (Swigut and Wysocka 2007)(De Santa et al. 2007). Temporal control seems important, H3K4me3 can deposited at promoters even when there is H3K27me3, setting up a poised chromatin configuration. Resolution of this bivalency by demethylation of H3K27me3 allow transcription elongation (Margueron and Reinberg 2011)(Liu et al. 2016). Current research suggests that KDM6B plays a more pivotal role in the brain cells than KDM6A and essential for neuronal development, synaptic plasticity, and cognitive function, with disruptions in its activity linked to cognitive impairments. In contrast, the role of KDM6A in brain function appears to be less prominent (Yang et al. 2017)(Wang et al. 2022a)(D’Oto et al. 2021)(Wang et al. 2022b). H3K27me3 is associated with repression of CD133, NESTIN, CD44 and SOX2 gene expression (Godfrey et al. 2020)(Boulland et al. 2012), are known for important stemness and pluripotency factors. KDM6B known to be a transcription activator epigenetically on CD44 and SOX2 gene promoters in different cancer cells (Shait Mohammed et al. 2022) and maintain pluripotency in hematopoietic stem cells in context dependent manner (Mallaney et al. 2019). KDM6B is reported as an important epigenetic modifier in maintenance and establishment of neuronal stem cells in dentate gyrus in mice (Gil et al. 2024). KDM6B reported to be recruited at transcription start site by interacting with different DNA binding proteins like T-Box TFs, Smad proteins, estrogen receptors and KLF4 (Akizu et al. 2010)(Miller et al. 2010)(Svotelis et al. 2011)(Huang et al. 2020). Another well-known transcription factor, p53 is reported to be a lineage determinant in RA induced differentiation by recruiting H3K27me demethylase KDM6B on target gene promoters (Akdemir et al. 2014)(Williams et al. 2014). P53 reported to physically associated with AT-hook protein like HMG1, p300 and TAFII250 for different cellular functions and nuclear localization (Imamura et al. 2001)(Jayaraman et al. 1998)(Wiederschain et al. 2003)(Allende-Vega et al. 2007), might have an accessory role in AKNA’s functions. Nodal signaling dependent developmental reprograming is H3K27me3 demethylation dependent which catalyzed by smad2/3 dependent KDM6B recruitment on targeted gene promoter (Dahle et al. 2010). There are other examples also where KDM6B either accumulated in the nucleus or recruited to the promoter by different nuclear proteins (Zhao et al. 2013)(Estarás et al. 2012).

As per available literature, prevailing view is that different TFs and nuclear proteins assists localization and functions of epigenetic modifiers like KDM6B and reprogram the gene expression pattern and depict cell fate. However, centrosomal functions and specifically transcription regulation by AKNA is poorly understood. Further investigation of how AKNA brings these two roles might focus further insights into cellular differentiation, immune response, and tumorigenesis. Herein, we have investigated how AKNA imparts as context-dependent molecular switch for chromatin function, in which condition AKNA retained in cytosol, how AKNA become mandatory for both stemness and differentiation. This report elaborates the functional role of AKNA in histone modifications, essential for activation of genes associated with stemness and differentiation in neuroblastoma lineages.

## Materials and methods

### Culture of Neuroblastoma cell lines

IMR32 and SH-SY5Y cell lines were procured from NCCS, Pune, India. These cell lines were grown in DMEM and F12 media respectively. 10% serum (Gibco -10270106) supplement and 1X anti-anti were added to the cultured media and incubated at 37°C in a 5% CO2 incubator. For induction of stemness, DMEM/F12 (1:1) was used instead of their cultured media for the cell lines. To treat the cells with different agents, cells were counted and seeded according to their rate of

### Drug treatments and MTT assay

To check the cytotoxicity of different treatments, cell viability was assessed using MTT (Himedia-TC191) in live cells. Cells were seeded at 10^3^ to 1.5*10^3^ per well in 96 well cell culture plate. After 24hrs of incubation in CO2 incubator, cells were treated with different concentrations of drugs for 48hrs because the downstream experiments need long exposure time. After that drug media was replaced with MTT media and incubated for 4-6 hrs followed by DMSO (Himedia – AS121) mediated solubilization of formazan crystals. The absorbance was captured at 570nm which was used to calculate cell viability as follows, % Viability =100 * mean OD (drug)/mean OD (control).

Drugs used in this study: Retinoic acid (Sigma-R2625), OAC1 (Santa Cruz-sc-397046). division. For microscopy experiments, 10-15% cells per well seeded to avoid overgrowth.

### Neurosphere formation assay

To determine the presence and prevalence of stem cells in cultured cells in different treatment, neurosphere formation assay was performed. Properly dissociated cells were seeded on a low attachment plate using DMEM/F12 media. Cultured were kept in CO2 incubator for 5-7 days. These cells easily form neurosphere and the images were taken using a phase contrast microscope (Olympus).

### In Vitro neuronal differentiation

IMR32 and SH-SY5Y cell lines were seeded at a density of 1×10^5^ cells per 6 well culture plate coated with poly L-lysine in normal growth media followed by overnight incubation for attachment. After 24hrs culture media was replaced by 2.5% FBS containing DMEM media with 10µM retinoic acid (RA) (Sigma-R2625-100MG) for SH-SY5Y and 5µM for IMR32. From the next day, half of the culture media from each well was replaced by fresh RA containing media for up to 5 days. Neurite outgrowth was assessed under a phase contrast microscope. Three different time points (1,3 &5^th^ day) differentiated cells were used for downstream experiments to study the gradual molecular changes during differentiation. Although different groups of researchers differentiate for variable time point according to their research interest. In this study we intended to induce differentiation for up to 5days to induce higher expression of differentiated neuronal markers and that was confirmed with expression of differentiation markers in downstream experiments (Kovalevich et al. 2021)(Xie et al. 2010).

### Colony formation assay

Singles cells were seeded in 35mm Petridish (500/well) and incubated for 24hrs, followed by replacement of the growth media respective treated media and incubated for another 10-15 days depending on cell line. Then the colonies were fixed with formaldehyde and stained with crystal violet. Excess stain was washed with tap water and image has been captured as described earlier (Kar and Patra 2018).

### siRNA and plasmid transfections

Transfection was performed using lipofectamine 3000 (Invitrogen L3000-15) in opti-MEM media following manufacturer’s instructions. Different concentration of siRNA (AKNA siRNA) was used and validated in western blotting for knockdown efficiency confirmation. For overexpression, 3-5 µg of plasmid (KDM6B) was used depending on the culture plate size and cell density.

### Apoptosis assay

Cells were seeded at 10000 per well on a coverslip inside a 6 well plate. Then after attachment, cells were treated with respective treatment and incubated for another 48hrs. After that, cells were washed with PBS and stained with acridine orange (100μg/mL) and EtBr (100μg/mL) for 5mins, followed by imaging under fluorescence microscope (Olympus).

### Caspase-3/7 Activity Assay

Protein extract from SH-SY5Y cells after treatment quantified and 20µg protein from each sample was incubated with Caspase-Glo® 3/7 (Promega, USA) for 2hr in dark at room temperature to check the cleavage of luminescent substrate. Luminescence was measured by luminometer and normalized with total protein content followed by data presented in a bar graph.

### RNA isolation and qRT-PCR

For gene expression quantification using qRT-PCR, RNA samples were isolated using TRIzol (Thermo - 15596018) method and phenol chloroform (himedia-MB063) precipitation. RNA samples were washed with 70% ethanol (Himedia-MB228) and after drying, dissolved in nuclease free water. After qualitative and quantitative analysis of RNA samples in agarose gel electrophoresis and nanodrop (Eppendorf), cDNA synthesis was performed using reverse transcriptase (Genesure – PGK163A) according to the manufacturer instructions. For qRT-PCR, 1µg of cDNA and 300nM respective gene primer were taken with SYBR-green PCR mix (Thermo – A25742) and performed qRT-PCR like previously described way (Kar et al. 2014). Data was analysis was performed according to Livak’s method (ΔΔCT calculation) using either GAPDH or ACTB as housekeeping gene.

### Nuclear and cytosolic protein fractionation

To obtain nuclear and cytosolic fractions, cells were incubated in low salt buffer (10mM HEPES/KOH pH-7.9, 10mM KCl, 1.5mM MgCl2, 1x Protease inhibitor cocktail) for 10mins at 4°C in a rotor. After centrifugation (2000g-5mins), supernatant was taken as cytoplasmic fraction. The remaining pellets were rewashed with low salt buffer for removing any residual cytoplasmic contamination. After that, remaining pellets were dissolved in high salt buffer (20 mM HEPES/KOH pH = 7.9; 420 mM NaCl; 1.5 mM MgCl2; 200 uM EDTA; 23% glycerol; 1xPIC) followed by incubation at 4°C for 20mins and centrifugation at 10,000rpm for 10mins. The

### Immunoblotting

Whole cell lysates were extracted using RIPA buffer (sigma –R0278-50ML) followed by centrifugation (12000rpm-15mins) and immunoblotting was performed by following the protocol adapted from (Manna et al. 2023a). In brief the protein samples were separated on 8-12% SDS-PAGE gels based on target protein molecular weight, resolved and transferred to nitrocellulose membrane (Axiva – 160300RI) using semidry transfer system (BioRad), blocked with 5% skim milk for 1hr at RT. All primary antibodies were diluted in 1% BSA (Himedia-MB083) dissolved in PBST, and the blots were incubated with primary antibodies at 4°C overnight. After removing the primary antibodies, blots were washed with PBST for three times (10mins*3), then they were probed with either anti-rabbit (Invitrogen-65-6120) or anti-mouse (Santa Cruz – SC516102) secondary antibody for 1hr at RT. Then, the blots were washed with PBST for three times (3*10mins), followed by visualization using ECL substrate (Thermo-34580) in a ChemiDoc instrument (BioRad).

### Immunocytochemistry

Immunoblotting was performed as per previous standard protocol (Sengupta et al. 2016). Cells were seeded on a coverslip in a 6-well plate, treated with different agents for different time point according to the requirements, followed by fixation using 4% formaldehyde for 5 minutes. Then the cells were permeabilized with PBST on ice for 10 minutes and 1hr incubation with 1% BSA for blocking. Then cells were incubated with respective primary antibodies overnight at 4°C in a humidified chamber. Next, the cells were washed with PBS (10min×3) and incubated with Alexa647/Alexa488 (anti-mouse - ab150119 or ant-rabbit - ab150077), followed by washing with PBS thrice and images were captured using Leica microsystems (STELLARIS 5).

### Coimmunoprecipitation

Whole cell lysates were used for co-immunoprecipitation assay. Protein samples (300ug) were incubated with respective primary antibodies for overnights at 4°C. Next day, each sample was poured with precleared A/G beads and incubated for 2hrs at 4°C. For negative control mouse IgG antibody was used. Then, immunoprecipitated complexes were washed with wash buffer for three times, followed by elution with 20ul Laemmli buffer and heating. Protein-protein interactions were analysed by western blotting.

### Chromatin Immunoprecipitation (ChIP) assay

Chromatin immunoprecipitation assay was performed using a kit method (Sigma-Imprint chromatin immunoprecipitation kit – CHP1) according to manufacturer’s instructions. Treated and control cells were fixed with 1% formaldehyde in PBS for 10 minutes in shaking condition. Formaldehyde neutralized by using 1.25M glycine for 5 minutes at RT. Centrifuged and washed with PBS and lysed (on ice) for 10 minutes in 300uL of ChIP lysis buffer, followed by sonication (0.3 mm probe sonicator) on ice for 10 minutes on 30s ON-30s OFF cycles at 30% efficiency and small fraction was checked for length of fragments. Then the samples were diluted by equal amount of ChIP dilution buffer. Rest of the protocol was as mentioned in the kit. The DNA samples were purified using PCR purification kit (Qiagen). % input was calculated according to standard method, as described earlier (Kirtana et al. 2023).

### Statistical analysis

Either student’s ‘t’ test or two-way ANOVA followed by Bonferroni correction were performed in Prism5 software to test the significance with sample number ≥3, and the test employed along with the p-values (* p < 0.05, ** p < 0.01, and *** p < 0.001. ns, no significance) and their inference is mentioned in figure legends.

## Results

### OAC1 induces pluripotency and stemness in neuroblastoma cells

In neuroblastoma cell lines (IMR32 and SH-SY5Y), OAC1 treatment led to significant increase in colony formation capacity and neurosphere generation, with a greater number and larger size of spheres (**Fig. 1A-B; Supp. Fig. S1**). These findings suggest that OAC1 enhances stem-like properties in these cells. To further investigate this, we assessed the expression of stemness and pluripotency markers. Western blot analysis revealed a time-dependent upregulation of CD133, CD44, OCT4, SOX2, and PAX6, with 24 to 48 hours of treatment being sufficient to induce robust expression of these markers (**Fig**. 1C-D). To evaluate whether OAC1 treatment represses neuronal differentiation, we performed immunostaining for the neuronal differentiation marker TUBB3, which showed a marked reduction in expression after 48 hours of treatment. In contrast, the expression of the neuroepithelial marker, Nestin was significantly increased. The upregulation of PAX6 and SOX2 was further confirmed by immunostaining, reinforcing the observation that OAC1 drives a transition toward a pluripotent, stem-like state in these neuroblastoma cell lines (**Fig.** 1E). These findings demonstrate that OAC1 treatment effectively reprograms neuroblastoma cells toward a stem-like state by inducing pluripotency genes and suppressing neuronal differentiation.

**Figure 1:**
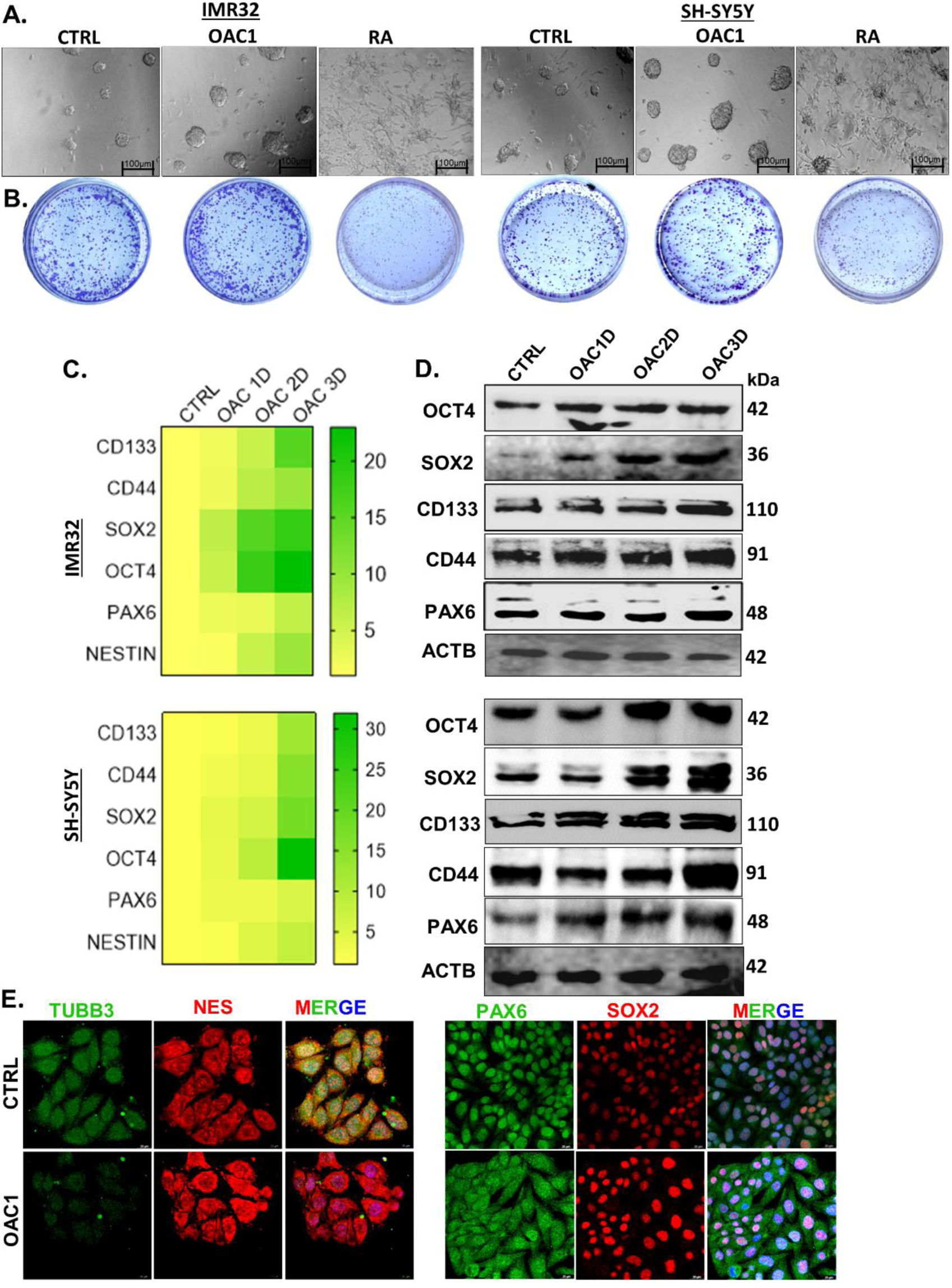
OAC1 efficiently induce stmeness and pluripotency in neuroblastoma cells. (A) Phase contrast microscopy of 3D neurosphere formation assay in OAC1 and RA treatment (9 Days) in IMR32 and SH-SY5Y cells. (B) Colony formation assay in IMR32 and SH-SY5Y cells treated with OAC1 and RA. (C) Heatmap represent qRT-PCR data in time dependent OAC1 treatment (1D-1 Day; 2D-2 Days; 3D-3 Days) in IMR32 and SH-SY5Y cell lines. (D) Immunoblotting analysis of stemness and pluripotency markers (CD133, CD44, OCT4, SOX2, PAX6) during time dependent OAC1 treatment in both cell lines; ACTB used as a loading control. (E) Immunocytochemistry images of different stemness and differentiation markers (NES, PAX6, SOX2, TUBB3) expression in OAC1 treatement. Images were acquired by laser confocal microscope at 63X under oil immerson (n>3, with at least 10 cells per focus area; Scale bar 20µm). All the experiments were performed three times or more (n≥3). Heatmap was generated using Prism 8 using statistically significant values and all other statistical analysis (either Anova or student ‘t’ test) was performed using Prism 5, (*)p < 0.05, (**) p < 0.01, and (***) p < 0.001. (ns), no significance.

### AKNA expression gradually increases during both OAC1, and retinoic acid induced cell fate transitions

Treatment of both IMR32 and SH-SY5Y cells with OAC1 is associated with multiple cells clustering and sphere-like forms, indicating the direction towards an enhanced stem-like state compared to control cells have more differentiated, attached morphology. Treatment with RA drives neuronal like differentiation with elongated, neurite-like extensions (**Fig.** 2A). There is a strong, time dependent increase in AKNA mRNA with OAC1 treatment (24h, 48h, 72h) in both cell lines, with a maximum at 72 hours. Conversely, RA treatment also increases AKNA, albeit less so than OAC1. AKNA protein expression reflects the mRNA findings, with a robust time dependent increase in OAC1-treated cells. RA treated cells also exhibit AKNA upregulation, although the increase is less than with OAC1. The quantified data validate that AKNA protein levels increase substantially with time with OAC1 treatment and, to a lesser extent, with RA. Maximum expression is seen at 72 hours of OAC1 treatment, whereas RA treatment is moderately induced (**Fig.** 2B-E). These finding suggests that OAC1 treatment induces strong expression of AKNA at the mRNA and protein levels, and this is concurrent with morphological changes characteristic of stemness. RA treatment induces upregulation of AKNA, but this is concurrent with morphological markers of neuronal differentiation. These indicates that AKNA can exert a context-dependent function in mediating stemness vs. differentiation in neuroblastoma cells.

**Figure 2:**
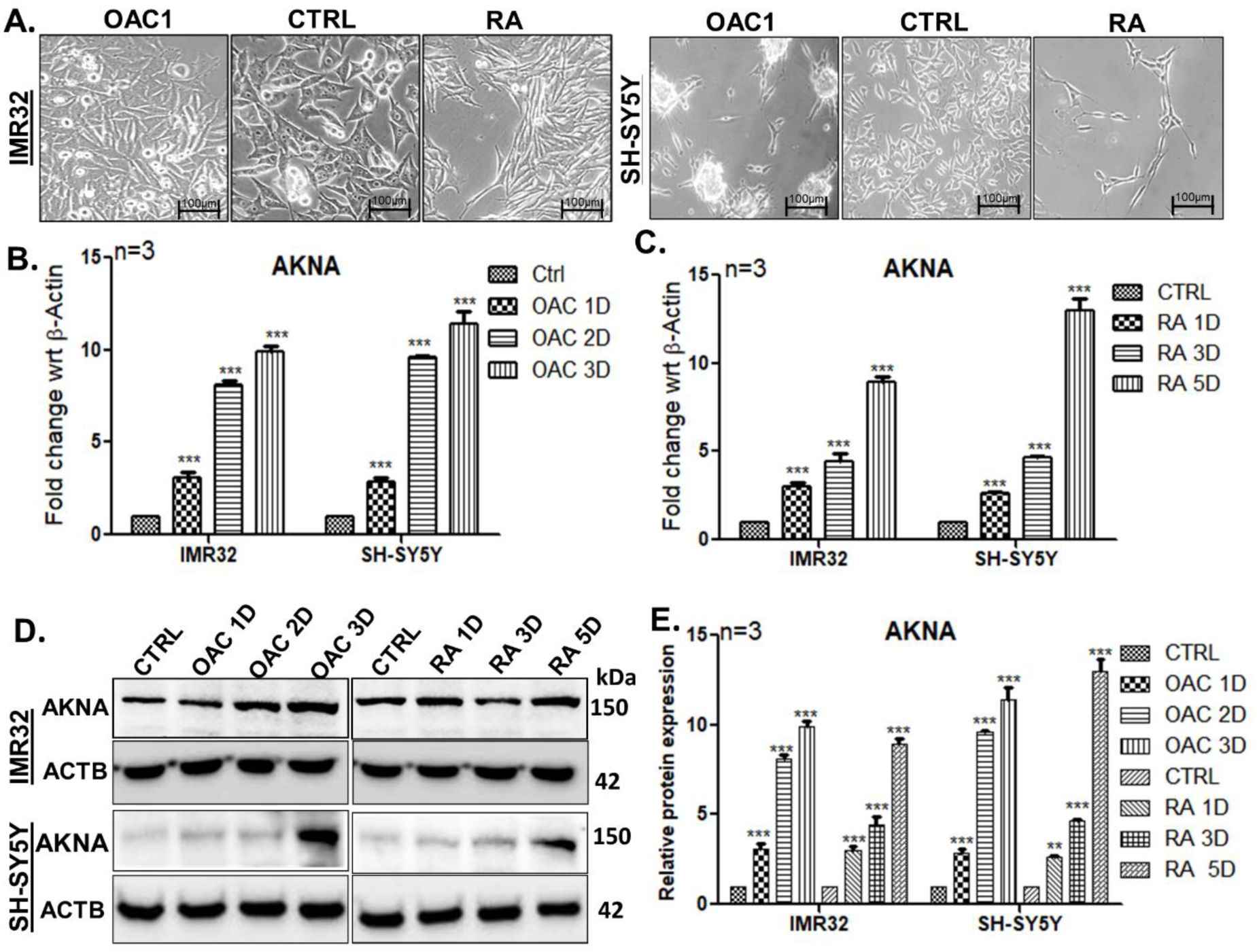
Upregulation of AKNA expression during both stemness and differentiation induction in neuroblastoma cells. (A) Morphological analysis in OAC1 and RA treatment in IMR32 and SH-SY5Y cells using microscopy. (B) qRT-PCR analysis for AKNA transcription rate during induction of stemness using OAC1in time dependent manner (1D-1 Day; 2D-2 Days; 3D-3 Days; 5D-5 Days) in both cell line. (C) AKNA transcription analysis in time dependent RA treatment. (D) Immunoblotting data of AKNA in OAC1 and RA treatment; beta actin (ACTB) taken as a loading control. (E) Bar graph represents relative protein expression visualized in immunoblots analysed by ImageJ software. All experiments were performed at least three times (n≥3) and statistical analysis (either Anova or student ‘t’ test) was performed using Prism 5, (*)p < 0.05, (**) p < 0.01, and (***) p < 0.001. (ns), no significance.

### In absence of AKNA, neuroblastoma cells failed to pass mitotic phase with impaired differentiation to mature neuronal cells

As the earlier data shown that OAC1 induces stemness and pluripotency whereas retinoic acid, known as differentiation inducer, both upregulates the expression of AKNA, so the effects of loss of functions of AKNA in stemness and differentiation was assessed. AKNA knockdown was performed using siRNA and knockdown efficiency was confirmed by western blotting (**Suppl. Fig. S2**). Next, in AKNA knockdown it was observed that stemness and pluripotency markers like OCT4, SOX2, CD133, CD44, NES, PAX6 expression were significantly downregulated. Differentiation markers like TUBB3, NEFM and NEFL were also downregulated in AKNA knockdown cells but intermediate progenitor and immature neuronal markers like TBR2 and TBR1 does not shows any significant changes (**Fig. 3A, D, Suppl. Fig. S3, S4**). These results also reflected in neurosphere and colony formation efficiency of the treated cells as a smaller number of colonies and smaller neurosphere were observed in AKNA KD cells (**Fig. 3B-C; Suppl. Fig. S5**). As the cells neither differentiating nor showing pluripotency, it prompts us to check the state of the cells. For that, cell cycle assay was performed and an increased G2/M phase arrest cells were observed in AKNA knockdown cells (**Fig. 3E**). To precisely known about AKNA effects on cell cycle, in AKNA knockdown cells chromosomal spreading was visualized using nuclear DNA specific dye (**Suppl. Fig. S6**). In AKNA knockdown it was visible from flow cytometry data that the cells which were arrested in G2/M phase were mostly arrested in mitotic stages, confirmed by immunostaining and immunoblotting with phase specific antibody (anti-H3S10P) (Fig. 3F-G). This cell cycle arrest leads to apoptotic cell death, determined by caspase glow assay and AO-EtBr staining. It is also evident that the apoptotic cell death is p53 dependent because p53 inhibition by Pifithrin-alpha (PFT-α) reduce the apoptosis **(Fig. 3H**; **Suppl. Fig. S7**). Physical association between AKNA and p53 (**Suppl. Fig**. **S8**) was observed, strengthen the p53 axis.

**Figure 3:**
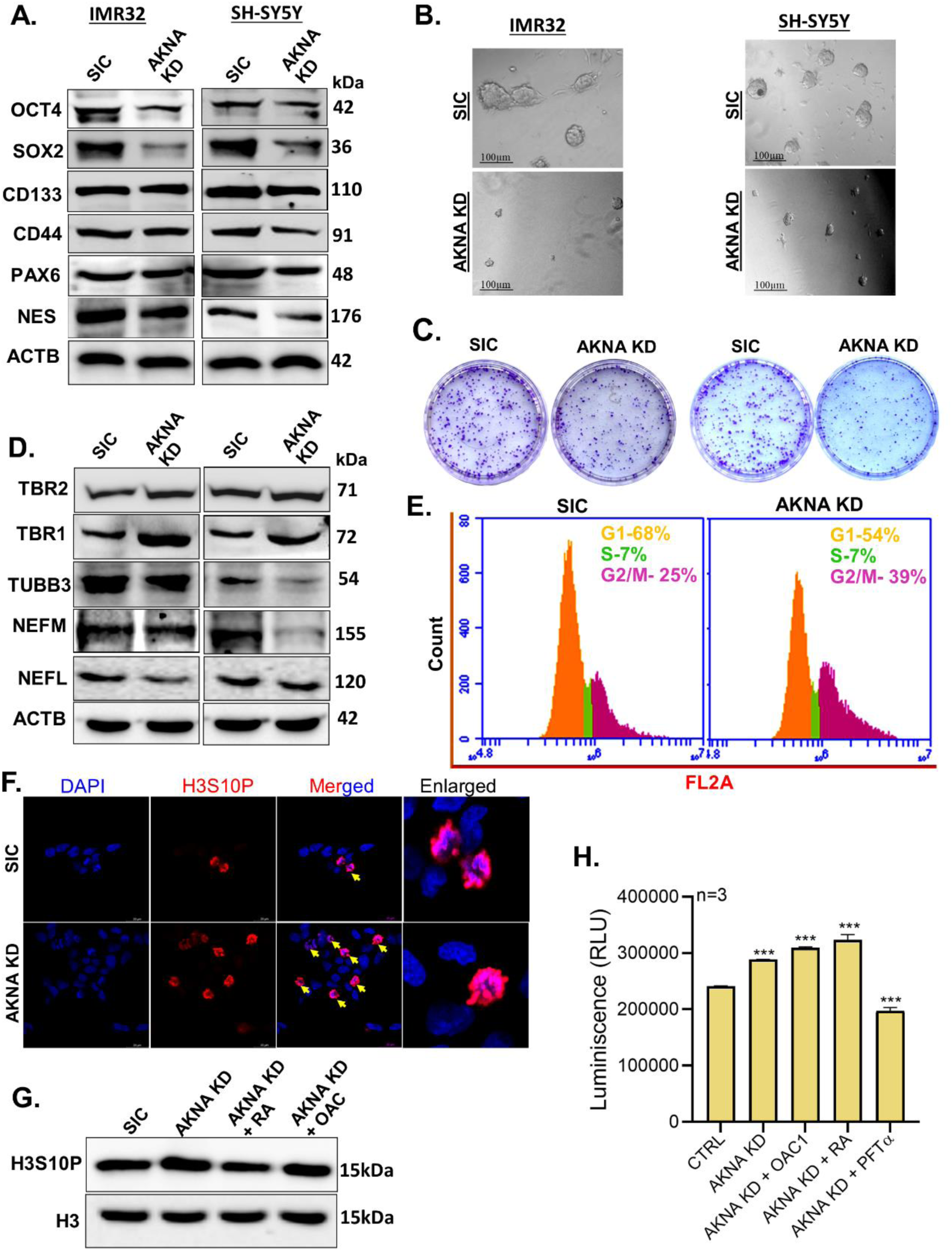
Absence of AKNA disrupts both stemness and terminal differentiation markers expression. (A) In AKNA knockdown stemness and pluripotency markers (CD133, CD44, OCT4, SOX2, NES and PAX6) protein expression was analysed by immunoblotting. (B) Neurosphere formation efficiency was checked in AKNA KD cells. (C) Colony formation efficiency of AKNA KD cells was visualized. (D) In AKNA KD, early and mature differentiation markers (TBR2, TBR1, TUBB3, NEFM, NEFL) protein expression were analysed in immunoblotting; ACTB used as a loading control. (E) Cell cycle analysis using flow cytometry in AKNA KD in IMR32 cells. (F) Immunostaining of H3S10p in AKNA KD IMR32 cells (G) Immunoblotting of H3S10P in AKNA knockdown; H3 used as loading control. (H) Caspase glow assay in AKNA KD with or without OAC1 and RA. (All the experiments were performed three times or more (n≥3) and statistical analysis (either Anova or student ‘t’ test) was performed using Prism 5, (*) p < 0.05, (**) p < 0.01, and (***) p < 0.001. (ns), no significance.

### AKNA is indispensable for determining both stemness and differentiation fate in neuroblastoma cells

Earlier results suggest that OAC1 induce pluripotency and stemness whereas RA induced differentiation, with increased expression of AKNA protein. To find out the role of AKNA in OAC1 induced stemness and RA induced differentiation, AKNA KD cells were treated with OAC1 and RA1. Interestingly it was observed that in absence of AKNA, OAC1 did not able to induce pluripotency and stemness in neuroblast cells, confirmed by immunoblotting. This was further reflected in the colony formation and neurosphere formation capacity of neuroblast cells. In absence of AKNA both neurosphere and colony formation capacity decreases even though the OAC1 present in the medium (**Fig. 4A-C, Suppl. Fig. S9**). In case of differentiation intermediate progenitor marker like TBR2 protein expression does not hamper in absence of AKNA, indeed slightly upregulated when RA is present. But postmitotic neuronal marker, TBR1 and mature neuronal markers like NEFL and NEFM expression were significantly decreased when AKNA is absent during RA mediated differentiation induction (**Fig. 4D**). In absence of AKNA, RA failed to differentiate neuroblastoma cells to fully matured neuronal cells and a greater number of bipolar morphological neuronal cells were observed (**Fig. 4E**). To nullify any off-target effects of chemically synthesized small molecule, OAC1 on stemness and pluripotency induction, a pluripotency inducing vector (CoMiP) was transfected independently and in absence of AKNA. In CoMiP treated cells pluripotency and stemness markers were significantly upregulated as expected but in absence of AKNA, it failed to induce pluripotency and stemness markers expression both at transcription and translational level in neuroblast cells (**Fig. 4F-G**). This result further strengthens the importance of AKNA in maintaining stemness in neuroblast cells.

**Figure 4:**
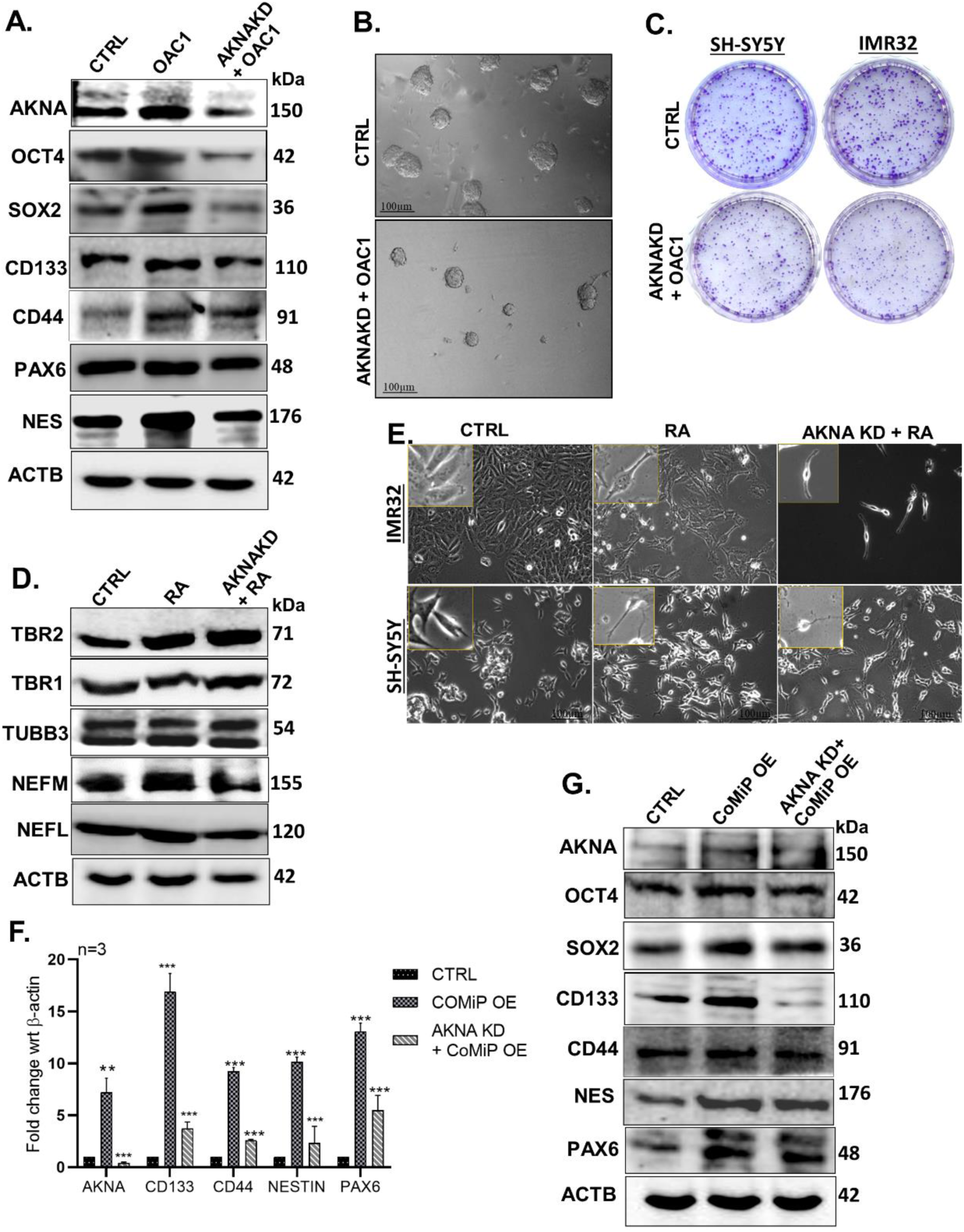
AKNA is indispensable during induction of stemness and differentiation. (A) Stemness markers (CD133, CD44, OCT4, SOX2, NES and PAX6) protein expressions were compared between only OAC1 treatment with double treatment of AKNA KD with OAC1 using immunoblotting; ACTB used as a loading control. (B) Neurosphere formation assay was performed and imaged in OAC1 treatment in absence of AKNA (OAC1+AKNAKD). (C) In same condition colony formation assay was performed. (D) Early and terminal differentiation markers (TBR2, TBR1, TUBB3, NEFM, NEFL) expression were analysed by immunoblotting; ACTB used as a loading control. (E) Differentiation morphology was visualized in same conditions under phase contrast microscopy (Scale bar 100µm). (F) Transcriptional rate of stemness markers in CoMiP treatment in presence or absence of AKNA were checked by qRT-PCR (CD133, CD44, NES and PAX6). (G) In same condition stemness markers translation were analysed by immunoblotting; ACTB used as loading control. All the experiments were performed at least three times (n≥3) and statistical analysis (either Anova or student ‘t’ test) was performed using Prism 5, (*) p < 0.05, (**) p < 0.01, and (***) p < 0.001. (ns), no significance.

### AKNA exhibits dynamic localization during stemness and differentiation induction and regulated by FAK signaling

To study the dual fate directional role of AKNA prompt us to study the cellular localization of AKNA protein in different treatment conditions. Surprisingly it was observed that OAC1 induce more nuclear localization of AKNA compared to the untreated cells, seen in the confocal microscopy and immunoblotting assay. Oppositely it has been found that RA treatment retains more AKNA protein outside the nucleus (**Fig. 5A-B**). Sodium orthovanadate (Tyrosine phosphatase inhibitor) treatment significantly increase AKNA’s cytosolic localization even in presence of OAC1 (**Fig. 5C-D**).

**Figure 5:**
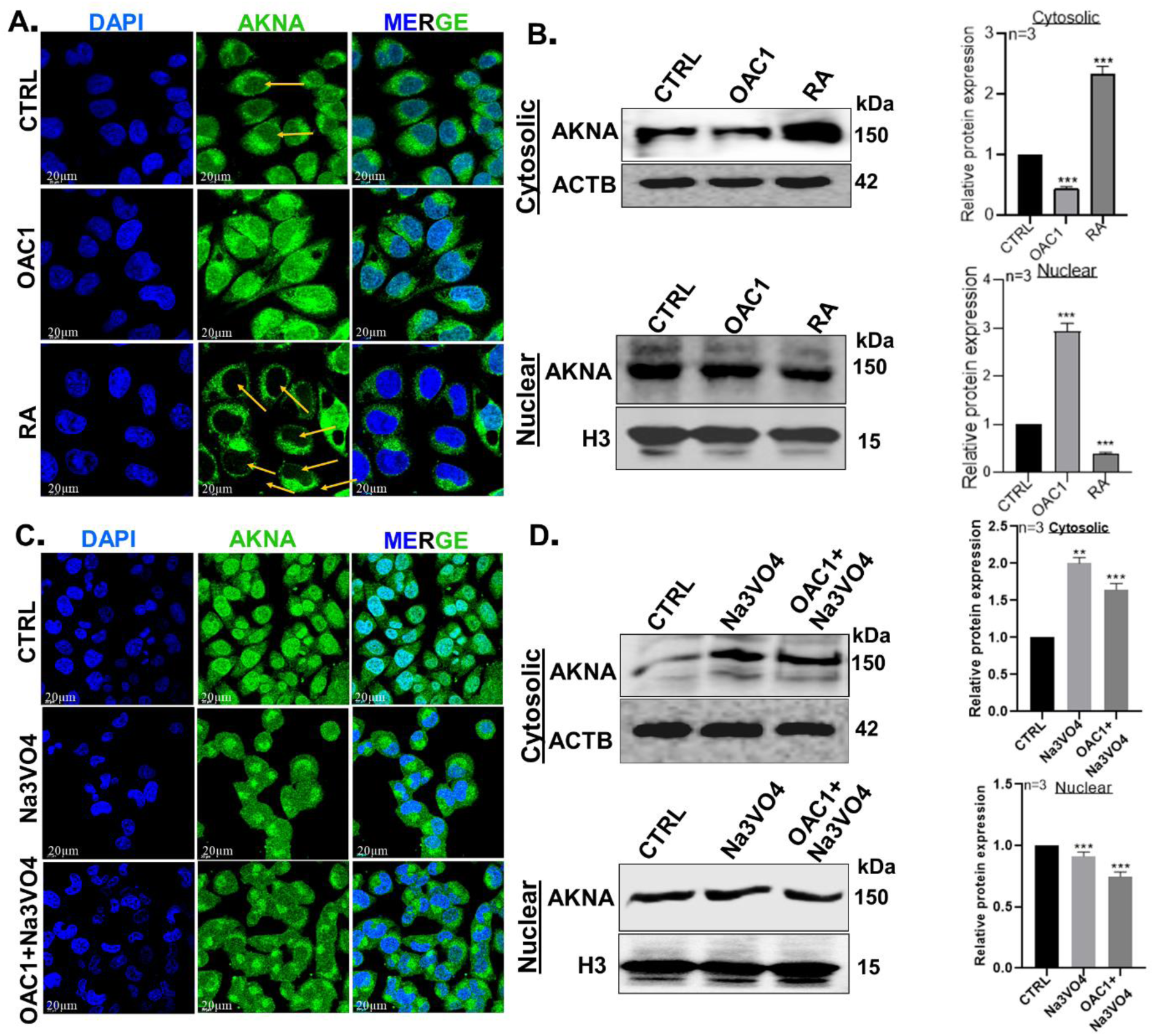
Cellular localization of AKNA during induction of fate transition is phosphorylation dependent and governed by FAK signaling. (A) AKNA localization was visualized in OAC1 and RA treatment (Green-AKNA; Blue-DAPI for DNA) in IMR32 cells. (B) Further validation of AKNA was determined by immunoblotting of AKNA using nuclear and cytosolic fraction; ACTB and H3 used as loading control for cytosolic and nuclear fraction respectively. (C) Localization of AKNA was visualized under oil immersion in Na3VO4 (a tyrosine phosphatase inhibitor) treatment with or without OAC in IMR32 cells, (With at least 10 cells per focus area; Scale Bar 20µm). (D) Expression of AKNA in cytosolic and nuclear fraction was analysed in the previous conditions. All the experiments were performed three times or more (n≥3) and statistical analysis (either Anova or student ‘t’ test) was performed using Prism 5, (*)p < 0.05, (**) p < 0.01, and (***) p < 0.001. (ns), no significance.

IMR32 cells treated with Na3VO4 for different timepoint showed that pFAK expression significantly upregulated while FAK expression almost similar to the earliest condition. To check the effects of FAK signaling on AKNA transcription, FAK signaling inhibited and activated by PF-573228 and FAK overexpression. In both conditions, AKNA expression remain unchanged (**Fig. 6A-B**). But in immunocytochemistry data, it was observed that FAK retain more AKNA in the cytosol whereas inhibition of FAK phosphorylation significantly increase AKNA inside the nucleus (**Fig. 6C-D**). FAK signaling inhibition also disrupts the RA mediated neuronal differentiation in neuroblastoma model (**Supp. Fig. S10**). This result suggests that AKNA’s dynamic localization is important for fate transition and regulated by FAK signaling.

**Figure 6:**
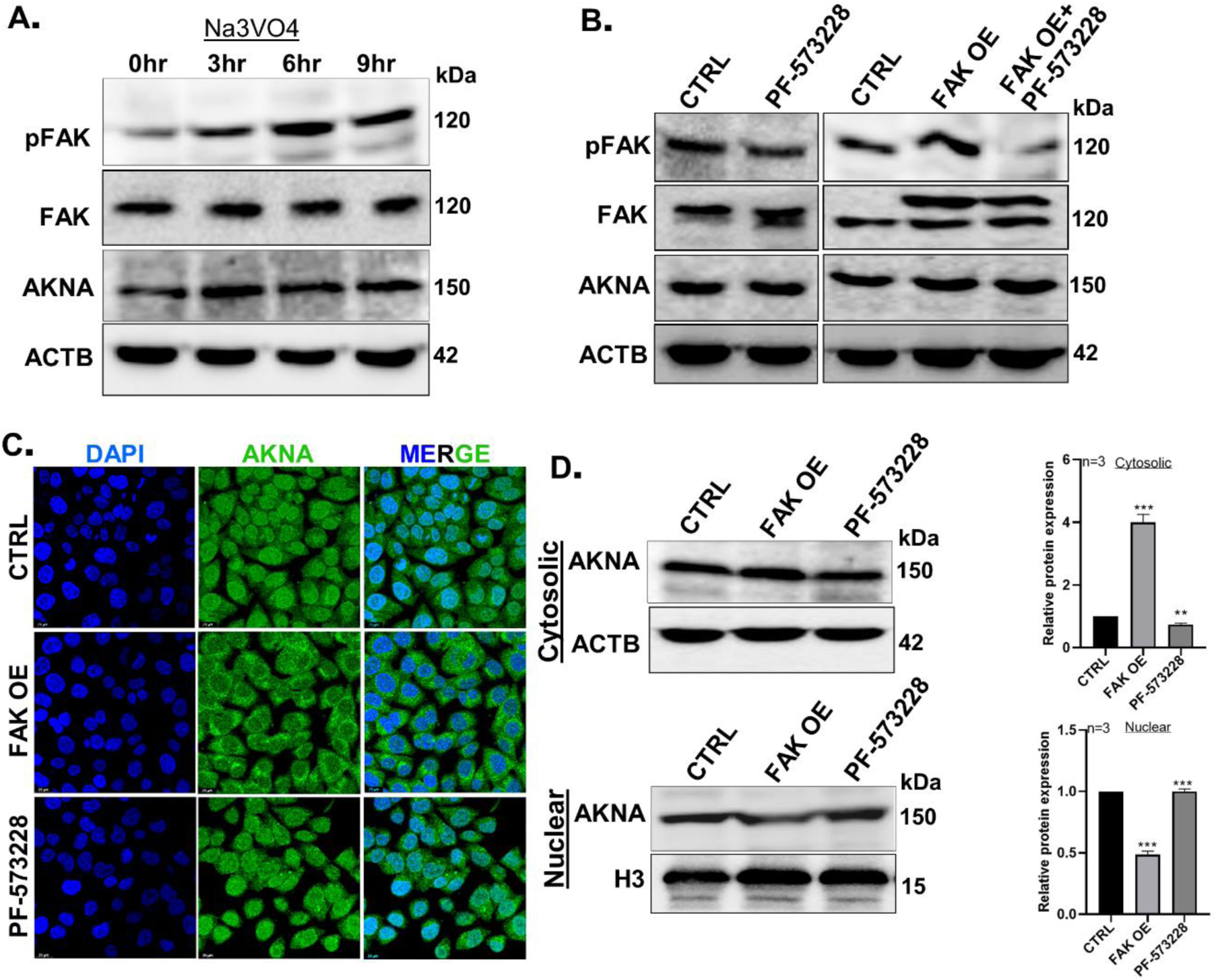
**(A)** Synchronized IMR32 cells treated with Na3VO4 for different time point (0hr, 3hrs, 6hrs and 9hrs) and pFAK, FAK and AKNA expression were analysed by immunoblotting. (B) FAK signaling was inhibited by PF-573228 and activated by FAK overexpression; immunoblotting was performed to check the effects on AKNA expression. (C) Role of FAK signaling in localization of AKNA was checked using confocal microscopy and (D) immunoblotting using nuclear and cytosolic fraction. All the experiments were performed three times or more (n≥3) and statistical analysis (either Anova or student ‘t’ test) was performed using Prism 5, (*)p < 0.05, (**) p < 0.01, and (***) p < 0.001. (ns), no significance.

### Accumulation of AKNA and reduction of H3K27me3 mark by KDM6B on gene promoters are associated with OAC1 induced pluripotency and stemness

To check the mechanisms of AKNA mediated transcriptional activation of stemness and pluripotency, ChIP assay was performed and observed that in OAC1 treatment occupancy of AKNA significantly increased on tested gene (NES, CD133, SOX2 and OCT4) promoter. In same condition transcriptional repressor mark H3K27me3 occupancy significantly decreased on the tested gene promoters. In time dependent treatment it was observed that H3K27me3 demethylase KDM6B upregulated gradually in both cell lines. Promoter occupancy of KDM6B also increased on tested gene promoters in OAC1 treatment (**Fig. 7A-D**). To verify the independent effects of KDM6B on pluripotency and stemness, knockdown and overexpression was performed followed by qRT-PCR and immunoblotting. In KDM6B KD significant reduction of stemness and pluripotency gene expression were observed in mRNA and protein level as well. Reverse results were observed in overexpression condition in both cell lines (**Fig. 7E-F; Suppl. Fig.S11**). It was also observed that KDM6B overexpression increase the neurosphere formation efficiency in both cell lines (**Fig. 7G**). ChIP experiment confirms that KDM6B upregulates the tested genes by removing repressive H3K27me3 from the gene promoters (**Fig. 7H**). This data suggests that AKNA possible associated with KDM6B to drive gene transcription in this tested model.

**Figure 7:**
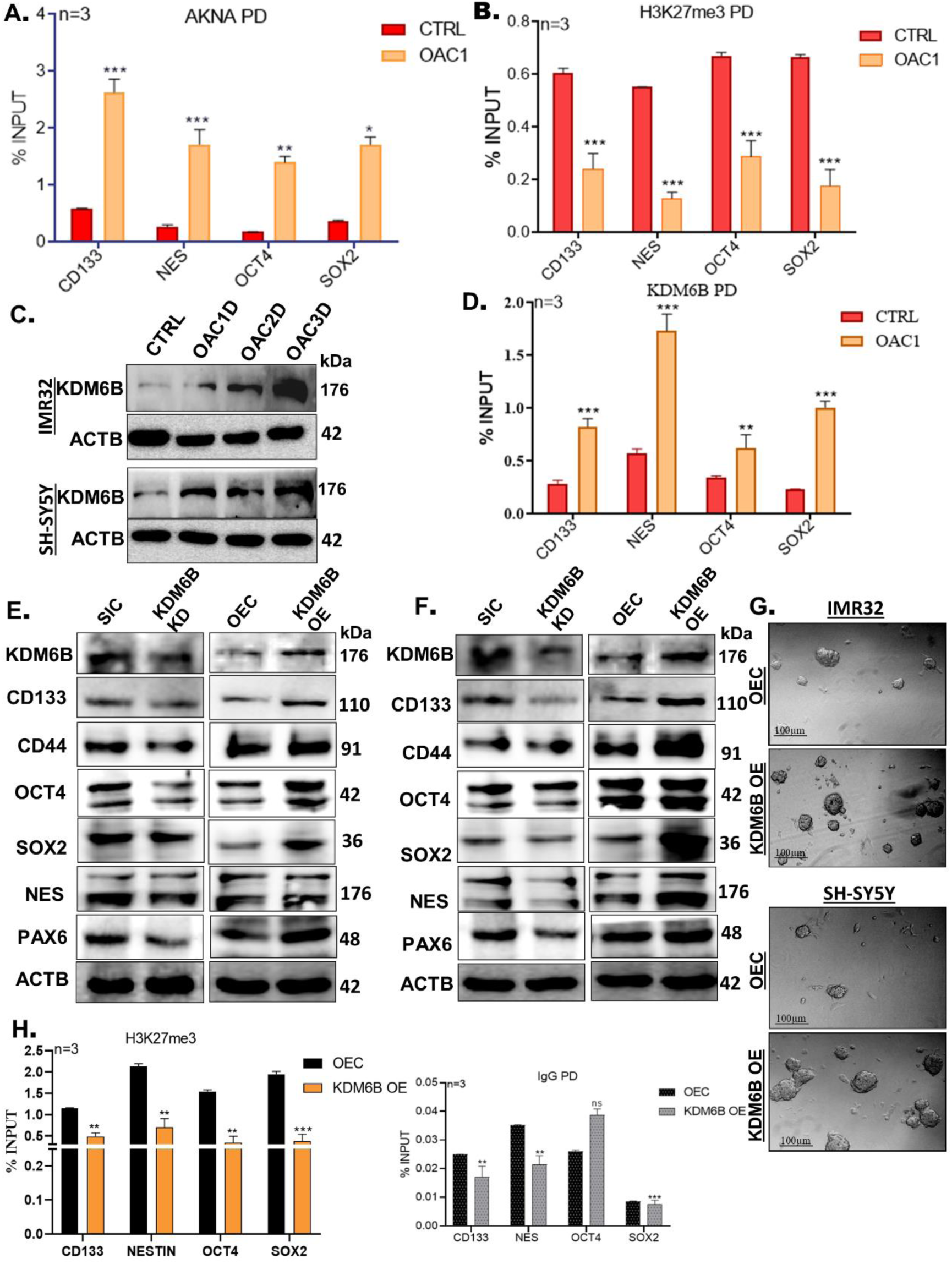
Transcriptional activation of pluripotency and stemness genes by AKNA is mediated by H3K27me3 demethylation and regulated by KDM6B. (A) Chromatin immunoprecipitation of AKNA during induction of stemness by OAC1 was analysed on CD133, NESTIN, SOX2 and OCT4 gene promoters. (B) H3K27me3 occupancy was checked in same conditions. (C) KDM6B expression analysis by immunoblotting in OAC1 and RA treatment in time dependent manner in both neuroblastoma cell lines; ACTB as loading control. (D) KDM6B occupancy was checked by ChIP assay in OAC1 treatment. (E) Stemness and pluripotency markers genes (CD133, CD44, OCT4, SOX2, NES and PAX6) expressions were analysed by western blotting in KDM6B knockdown and overexpression conditions; ACTB used as loading control. (F) Neurosphere formation assay was performed in KDM6B overexpression condition and images were captured by microscopy. (F) Chromatin immunoprecipitation assay was performed to check the occupancy of H3K27me3 on studied gene promoters along with negative control IgG. All the experiments were performed three times or more (n≥3) and statistical analysis (either Anova or student ‘t’ test) was performed using Prism 5, (*)p < 0.05, (**) p < 0.01, and (***) p < 0.001. (ns), no significance.

### AKNA is essential for KDM6B mediated H3K27me3 demethylation and subsequent acetylation of H3K27 to H3K27ac for gene expression

To check the interdependency, KDM6B overexpressed in AKNA knockdown cells and observed that in absence of AKNA, transcription of the panel of genes, including CD133, CD44, NES, OCT4, SOX2 and PAX6 were failed, which otherwise were upregulated by KDM6B. It was further validated by immunoblotting (**Fig. 8A-B**). Neurosphere formation capacity also compromised for KDM6B overexpressed cells in absence of AKNA (**Fig. 8C)**. To find out the possible mechanisms of AKNA dependency of KDM6B, ChIP assay was performed. It was clearly observed that when AKNA is less occupied on the tested gene promoters, KDM6B occupancy also decreased (**Fig. 8D-E**). This data confirmed that H3K27me3 demethylase KDM6B heavily rely upon AKNA for transcription of pluripotency and stemness related genes in neuroblastoma cells.

**Figure 8:**
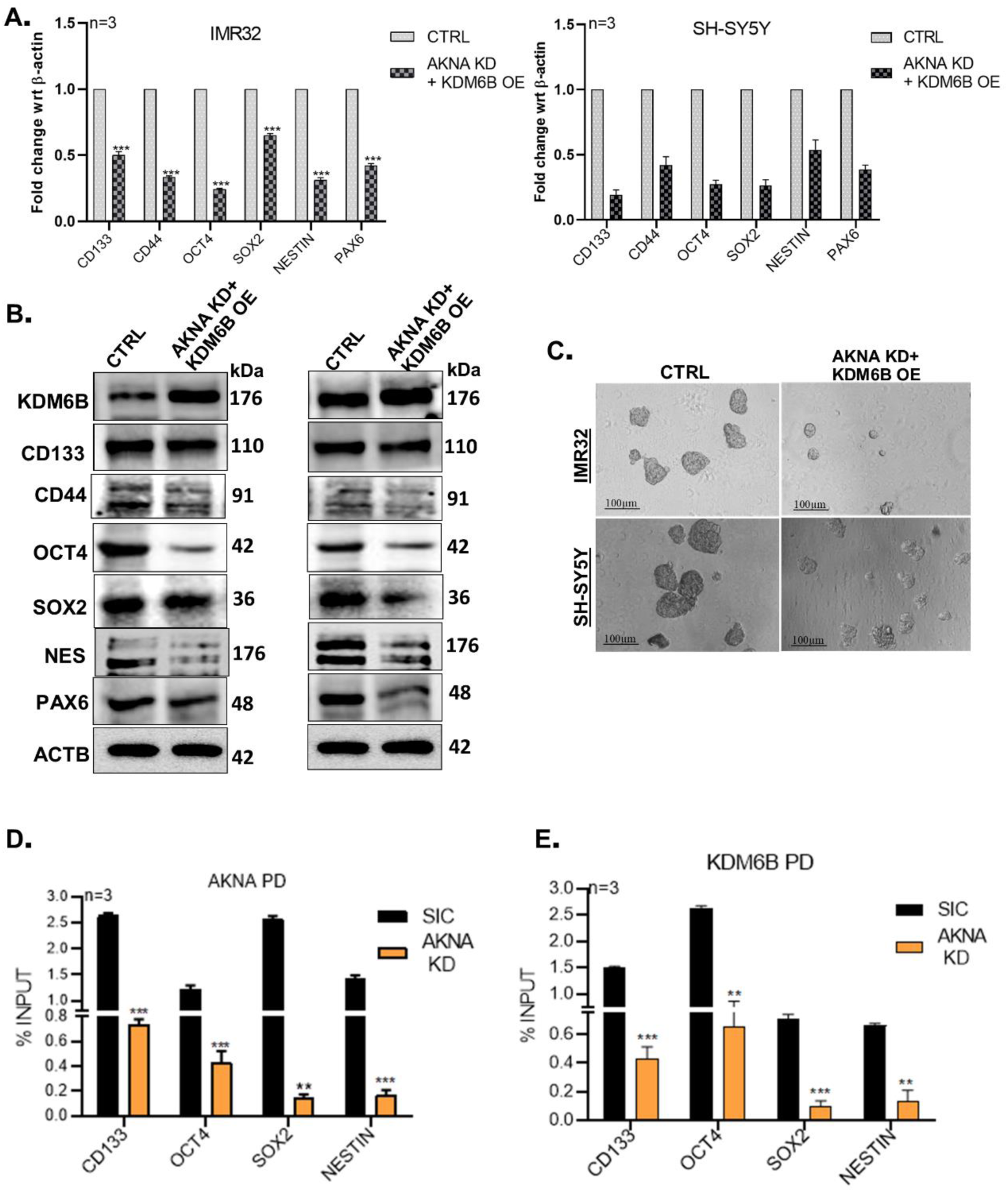
KDM6B mediated activation of pluripotency and stemness is AKNA dependent. (A) Transcription of stemness and pluripotency markers genes (CD133, CD44, OCT4, SOX2, NES and PAX6) were analysed by qRT-PCR in double treatment of AKNA KD and KDM6B OE in both IMR32 and SH-SY5Y cells. (B) That was further validated by immunoblotting, ACTB used as loading control. (C) Neurosphere formation assay was performed in previous double treatment conditions. (D) Chromatin immunoprecipitation was performed to check the change in occupancy of AKNA in AKNA KD cells. (E) In same condition KDM6B occupancy was analysed by ChIP assay. All the experiments were performed at least three times (n≥3) and statistical analysis (either Anova or student ‘t’ test) was performed using Prism 5, (*) p < 0.05, (**) p < 0.01, and (***) p < 0.001. (ns), no significance.

### AKNA and KDM6B interaction is indispensable for OAC1 driven stemness factors and pluripotency associated gene expression

To further confirm the effects of interdependency between AKNA and KDM6B, ChIP experiment was performed. It was observed that H3K27me3 is high in AKNA KD and when AKNA is absent KDM6B overexpression is not sufficient to demethylates H3K27me3 from the tested gene promoters. The mechanism behind such finding was that when AKNA absent, KDM6B failed to occupy the gene promoters as compared to the KDM6B overexpression only (**Fig. 9A-B**). Further it was also traced that H3K27 acetyl transferase p300/CBP complex along with H3K27ac depositions was higher in KDM6B overexpressed cells but decreased the occupancy when AKNA was knocked down in cells, irrespective of normal or ectopic overexpression of KDM6B (**Fig. 9C, Suppl. Fig.S12**). KDM6B physically interacts with different TFs to activate gene transcription by removing methylation from H3K27me3 and from above observation in this study its dependency on AKNA evoked to study their physical association. From immunocytochemistry data it was apparent that AKNA and KDM6B colocalize in the neuroblast cells and further immunoblotting data confirmed that AKNA protein physically interacts with KDM6B in different treatment conditions. Although started with equal amount of whole cell protein samples but the magnitude of interaction between KDM6B and AKNA is high in stemness induction but need further investigations to confirm (**Fig. 9D-E**). So, AKNA possibly promote the occupancy of KDM6B on gene promoters during induction of stemness in neuroblastoma cells.

**Figure 9:**
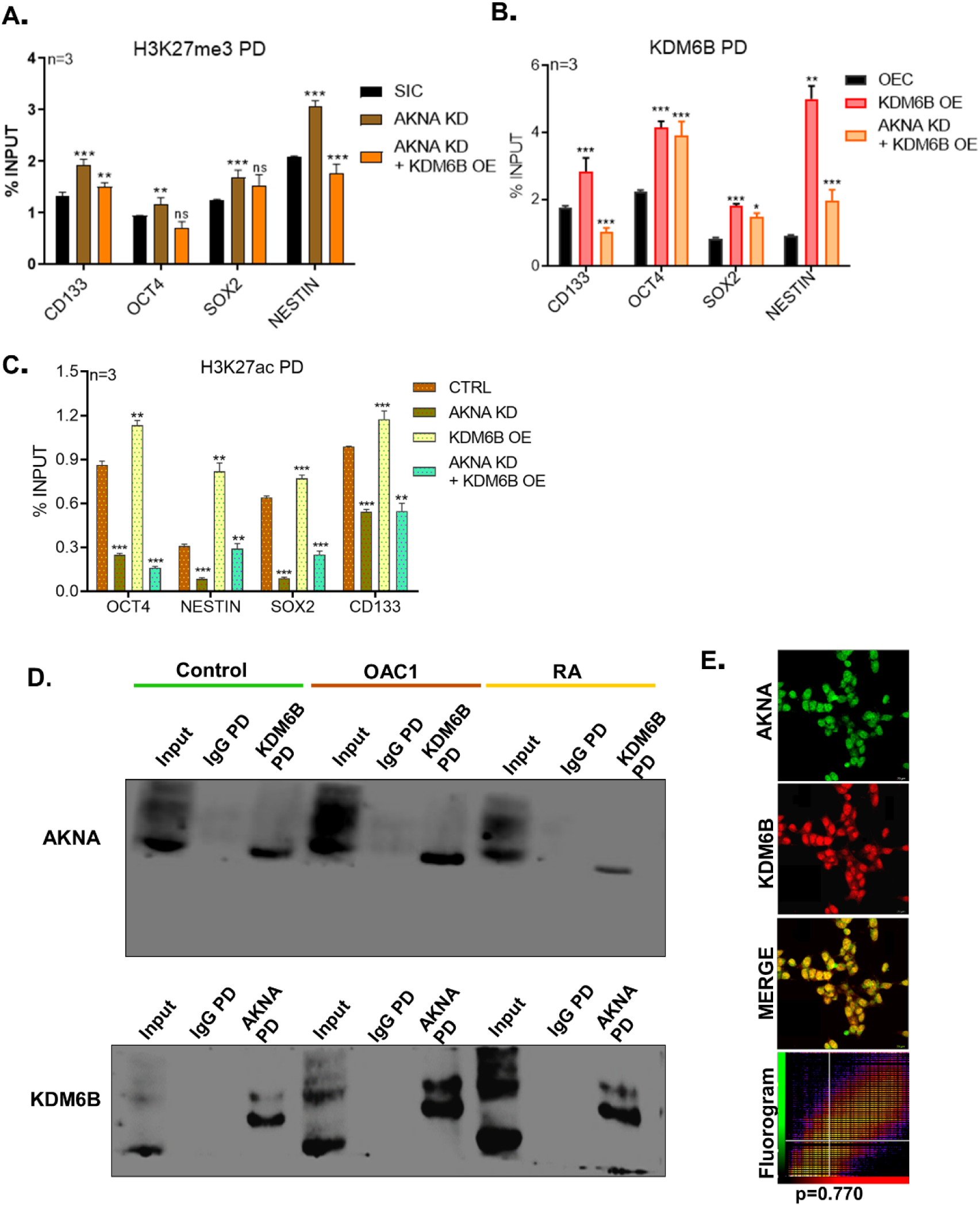
Promoter occupancy of KDM6B on tested genes is AKNA dependent by physical association. (A) H3K27me3 status on tested gene (CD133, NES, OCT4, SOX2) promoters were analysed by ChIP assay in AKNA KD and KDM6B OE in absence of AKNA. (B) In similar condition H3K27me3 demethylase KDM6B occupancy was checked by ChIP assay. (C) H3K27ac status was studied in similar conditions. (D) There interdependency was further validated by Coimmunoprecipitation experiment using whole cell extracts from control, OAC1 and RA treated IMR32 cells, followed by immunoprecipitation either by AKNA and KDM6B and immunoblotting with opposite antibody. Mouse IgG was used as a negative control. (E) This physical association was further confirmed by immunoblotting using both anti-AKNA and anti-KDM6B antibodies and a fluorogram was prepared (p>0.5) to check the associations. All the experiments were performed at least three times (n≥3) and statistical analysis (either Anova or student ‘t’ test) was performed using Prism 5, (*) p < 0.05, (**) p < 0.01, and (***) p < 0.001. (ns), no significance.

## Discussion

This study demonstrated that AKNA function as a key regulator for neuronal lineage fate determinant in neuroblast cell model in a context dependent manner. IMR32 and SH-SY5Y cells treated with OAC1 reprogrammed into enhanced pluripotent stemlike cells with enhanced self renewal and proliferation capacity, evident from colony and neurosphere formation study which was further confirmed by upregulated expression of markers like, OCT4, SOX2, NES, CD133, CD44 and PAX6. Simultaneously, OAC1 suppressed the differentiation, confirmed by reduction in expression of TUBB3, a well-known differentiation marker (Guo et al. 2010). This results aligned with the established theory of small molecules or factors capacity to reprogram fate determined cells into pluripotent cells (Li et al. 2012)(Takahashi and Yamanaka 2006)(Yu et al. 2007). In opposite directions, retinoic acid (RA) induce neuronal differentiation characterized by neurites like extension and apparent morphological changes (Kovalevich et al. 2021)(Encinas et al. 2000) (**Fig. 1-2**). Since the expression of AKNA was upregulated in both stemness and differentiation, demonstrating its dual impact in cellular and molecular context.

Depletion of AKNA arrest the cell cycle in mitotic phase which signifies that its role in cell cycle regulation may be linked with its role in fate determination. This mitotic arrest aligned with the previous findings of AKNA’s role in CD40 and CD40L regulation in lymphocytes maturation and it’s important role in M phase transition (Siddiqa et al. 2001)(Ramírez-González et al. 2023)(Lange et al. 2009). The shortfall of OCT4-SOX2-KLF4-cMYC tetrad (CoMiP plasmid) to induce stemness in absence of AKNA make it an unique TF from other TFs, like LIN28 and NANOG specifically in neuronal lineage (Yu et al. 2007)(Chambers et al. 2003) (**Fig. 3-4**). This dual functions of AKNA on fate and proliferation warrants further studies, plausibly through single cell RNA sequencing to decipher the dynamics of AKNA in fate decision (Trapnell et al. 2014).

On the contrary, in RA mediated differentiation retained AKNA predominantly in cytosol, correlated with the expression of neuronal differentiation markers and neurites formation (Harasym et al. 2017)(Tan et al. 2015). Earlier report suggests Ser/Thr phosphatase inhibition shifts the centrosomal localization of AKNA (Camargo Ortega et al. 2019) and here in this study it was observed that Tyrosine phosphatase inhibition retained AKNA in the cytosol. The mechanisms of tyrosine phosphorylation in localization of AKNA need further studies. Retinoic acid known to activate FAK signaling (Sanchez et al. 2016) and described for its importance in neurodevelopmental processes (Navarro and Rico 2014). FAK signaling here found to be regulates the localization of AKNA possibly by phosphorylation by an unknown mechanism. This dynamic shifts of AKNA in cellular compartment for fate determination are very much similar to the other TFs like SOX2, depending on the cellular context and interactions (Wang et al. 2012)(Schaefer et al. 2024). In the absence of AKNA, disruption of neuronal differentiation (e.g., TUBB3, NEFL, NEFM downregulated) without affecting intermediate progenitors cell identity (e.g., TBR2 unchanged) and generation of bipolar neuron like cells in presence of RA signifies that cytosolic AKNA possibly govern the later stages of neuronal differentiation, possibly post mitotic neuronal cells as mitotic phase arrest observed in knockdown condition. Such stage specific role are well known in cortical neurogenesis where different TFs orchestrate the process in a sequential manner (Greig et al. 2013) (Hsieh)(**Fig. 4-6**).

The OAC1 induced stem like states reflects the transcriptional landscape of embryonic stem cells, in which OCT4, SOX2 and PAX6 forms a pluripotency network hub (Yeo and Ng 2012)(Young 2011)(Sansom et al. 2009). This study demonstrated that in a time dependent manner these pluripotency markers were upregulated with simultaneously nuclear accumulation of AKNA protein, which sites AKNA as a possible transcriptional regulator in neuroblast stemness. This is reinforced by the physical interaction between AKNA and KDM6B, histone H3K27me3 demethylase important for activating pluripotency and stemness by removing methylation mark (Ding et al. 2021)(Akiyama et al. 2016). The interdependency between KDM6B and AKNA illustrated by the insufficiency of KDM6B to maintain stemness in absence of AKNA, which reflects the molecular basis of neural stem cell where epigenetic modifiers associated with TFs to established cell fates (Mirizio et al. 2025)(Matsui et al. 2024)(Taberlay et al. 2011). Strikingly in this study AKNA knockdown abolish the efficiency of KDM6B to occupy the promoters of pluripotency and stemness genes (OCT4, SOX2, CD133, NES), evidenced by ChIP data. This is allied with the role of KDM6B in neurodevelopment (Liu and Xiao 2011)(Stolerman et al. 2019)(Agger et al. 2007) (**Fig**. **7**).

Interaction between AKNA and KDM6B delivered a mechanistic axis link between epigenetic modification with transcriptional control on neuroblast plasticity. Induction of stemness and pluripotency upregulates KDM6B during OAC1 treatment that reduces repressive H3K27me3 on promoters which are aligned with its prevalent functions in neuronal stem cell maintenance (Gil et al. 2024)(Shait Mohammed et al. 2022)(Park et al. 2014). In absence of AKNA, occupancy of KDM6B are decreased in KDM6B overexpression conditions suggests that AKNA promotes and maybe stabilized the binding of KDM6B, as founds in other TFs and demethylase complex (De Santa et al. 2007).

These results will reinforce the understanding about neuronal development and differentiation process. Retinoic acid drives neuronal differentiation, is a recognized model for studying neuronal lineage commitment for embryogenesis (Maden 2007), whereas OAC1 drives stemness by AKNA, advocate dual regulating mechanisms. However, the context dependent role of AKNA entangled the mechanism and prerequisites upstream regulation study, like FAK signaling possibly through post translational modifications (Filtz et al. 2013) of AKNA, and its functions in neuronal lineage commitment (Jessell 2000).

In conclusion, AKNA emerges as a molecular keystone for neuronal lineage commitment by integration of transcription factor with epigenetic cues through an association with KDM6B. This work demonstrated a novel regulatory horizon that guide the balance between stemness and differentiation, presenting a foundation for orchestration of neurodevelopmental plasticity with broadscale implication in developmental biology and regenerative medicines.

## Acknowledgement

S. Manna and A. Roy are thankful to NIT-Rourkela for fellowships under the Institute Research Scheme, NIT-Rourkela. R. Kirtana received fellowship from CSIR, Govt. of India. Niharika was the JRF/SRF in the SERB project (project No.: EMR/2016/007034) and later received fellowship under the Institute Research Scheme, NIT Rourkela. Dr. P. Dash is the project research scientist-1 in the ICMR project (project No.: IIRP-2023-2134). T. Baral, J. Mishra, S. Chakraborty, P. Nandi, and P. Mishra are thankful to UGC for fellowships, Govt. of India. B. Pradhan is receiving fellowship from CSIR, Govt. of India.

## Author contribution

S. K. P., Conceived, Conceptualize and funding; S. M., Conceptualize, Methodology, **S**oftware, collection & curation of data and drafting of the manuscript. K. R., Methodology, collection & curation of data; T. B., J. M., P. N., S. C., N. M., A. R., P. M., B. P. & P. D. edited data and manuscript proofreading; S. K. P. edited the draft manuscript to its final form.

## Data availability

No datasets were generated or analyzed in this study.

## References

Agger K, Cloos PAC, Christensen J, Pasini D, Rose S, Rappsilber J, Issaeva I, Canaani E, Salcini AE, Helin K. 2007. UTX and JMJD3 are histone H3K27 demethylases involved in HOX gene regulation and development. Nature 449: 731–734.

Akdemir KC, Jain AK, Allton K, Aronow B, Xu X, Cooney AJ, Li W, Barton MC. 2014. Genome-wide profiling reveals stimulus-specific functions of p53 during differentiation and DNA damage of human embryonic stem cells. Nucleic Acids Res 42: 205–223. https://pubmed.ncbi.nlm.nih.gov/24078252/ (Accessed March 12, 2025).

Akiyama T, Wakabayashi S, Soma A, Sato S, Nakatake Y, Oda M, Murakami M, Sakota M, Chikazawa-Nohtomi N, Ko SBH, et al. 2016. Transient ectopic expression of the histone demethylase JMJD3 accelerates the differentiation of human pluripotent stem cells. Dev 143: 3674–3685. https://pmc.ncbi.nlm.nih.gov/articles/PMC5087640/ (Accessed March 27, 2025).

Akizu N, Estarás C, Guerrero L, Martí E, Martínez-Balbás MA. 2010. H3K27me3 regulates BMP activity in developing spinal cord. Development 137: 2915–2925. https://pubmed.ncbi.nlm.nih.gov/20667911/ (Accessed April 1, 2025).

Allende-Vega N, Saville MK, Meek DW. 2007. Transcription factor TAFII250 promotes Mdm2-dependent turnover of p53. Oncogene 26: 4234. https://pmc.ncbi.nlm.nih.gov/articles/PMC2695134/ (Accessed March 25, 2025).

Alver BH, Kim KH, Lu P, Wang X, Manchester HE, Wang W, Haswell JR, Park PJ, Roberts CWM. 2017. The SWI/SNF chromatin remodelling complex is required for maintenance of lineage specific enhancers. Nat Commun 2017 81 8: 1–10. https://www.nature.com/articles/ncomms14648 (Accessed March 25, 2025).

Aravind L, Landsman D. 1998. AT-hook motifs identified in a wide variety of DNA-binding proteins. Nucleic Acids Res 26: 4413–4421. 10.1093/nar/26.19.4413 (Accessed March 25, 2025).

Boulland JL, Mastrangelopoulou M, Boquest AC, Jakobsen R, Noer A, Glover JC, Collas P. 2012. Epigenetic Regulation of Nestin Expression During Neurogenic Differentiation of Adipose Tissue Stem Cells. https://home.liebertpub.com/scd 22: 1042–1052. https://www.liebertpub.com/doi/10.1089/scd.2012.0560 (Accessed March 25, 2025).

Camargo Ortega G, Falk S, Johansson PA, Peyre E, Broix L, Sahu SK, Hirst W, Schlichthaerle T, De Juan Romero C, Draganova K, et al. 2019. The centrosome protein AKNA regulates neurogenesis via microtubule organization. Nature 567: 113–117. https://pubmed.ncbi.nlm.nih.gov/30787442/ (Accessed March 12, 2025).

Chambers I, Colby D, Robertson M, Nichols J, Lee S, Tweedie S, Smith A. 2003. Functional expression cloning of Nanog, a pluripotency sustaining factor in embryonic stem cells. Cell 113: 643–655. https://pubmed.ncbi.nlm.nih.gov/12787505/ (Accessed March 30, 2025).

Cheung YT, Lau WKW, Yu MS, Lai CSW, Yeung SC, So KF, Chang RCC. 2009. Effects of all-trans-retinoic acid on human SH-SY5Y neuroblastoma as in vitro model in neurotoxicity research. Neurotoxicology 30: 127–135. https://pubmed.ncbi.nlm.nih.gov/19056420/ (Accessed March 12, 2025).

Collins BE, Greer CB, Coleman BC, Sweatt JD. 2019. Histone H3 lysine K4 methylation and its role in learning and memory 06 Biological Sciences 0604 Genetics 11 Medical and Health Sciences 1109 Neurosciences. Epigenetics and Chromatin 12: 1–16. 10.1186/s13072-018-0251-8.

D’Oto A, Fang J, Jin H, Xu B, Singh S, Mullasseril A, Jones V, Abu-Zaid A, von Buttlar X, Cooke B, et al. 2021. KDM6B promotes activation of the oncogenic CDK4/6-pRB-E2F pathway by maintaining enhancer activity in MYCN-amplified neuroblastoma. Nat Commun 2021 121 12: 1–19. https://www.nature.com/articles/s41467-021-27502-2 (Accessed March 12, 2025).

Dahle Ø, Kumar A, Kuehn MR. 2010. Nodal signaling recruits the histone demethylase Jmjd3 to counteract polycomb-mediated repression at target genes. Sci Signal 3. https://pubmed.ncbi.nlm.nih.gov/20571128/ (Accessed March 12, 2025).

De Santa F, Totaro MG, Prosperini E, Notarbartolo S, Testa G, Natoli G. 2007. The histone H3 lysine-27 demethylase Jmjd3 links inflammation to inhibition of polycomb-mediated gene silencing. Cell 130: 1083–1094. https://pubmed.ncbi.nlm.nih.gov/17825402/ (Accessed March 12, 2025).

Ding Y, Yao Y, Gong X, Zhuo Q, Chen J, Tian M, Farzaneh M. 2021. JMJD3: a critical epigenetic regulator in stem cell fate. Cell Commun Signal 19: 1–9. https://biosignaling.biomedcentral.com/articles/10.1186/s12964-021-00753-8 (Accessed March 27, 2025).

Dorafshan E, Kahn TG, Glotov A, Savitsky M, Walther M, Reuter G, Schwartz YB. 2019. Ash1 counteracts Polycomb repression independent of histone H3 lysine 36 methylation. EMBO Rep 20. https://pubmed.ncbi.nlm.nih.gov/30833342/ (Accessed March 25, 2025).

Encinas M, Iglesias M, Liu Y, Wang H, Muhaisen A, Ceña V, Gallego C, Comella JX. 2000. Sequential treatment of SH-SY5Y cells with retinoic acid and brain-derived neurotrophic factor gives rise to fully differentiated, neurotrophic factor-dependent, human neuron-like cells. J Neurochem 75: 991–1003. https://pubmed.ncbi.nlm.nih.gov/10936180/ (Accessed March 27, 2025).

Estarás C, Akizu N, García A, Beltrán S, de la Cruz X, Martínez-Balbás MA. 2012. Genome-wide analysis reveals that Smad3 and JMJD3 HDM co-activate the neural developmental program. Development 139: 2681–2691. https://pubmed.ncbi.nlm.nih.gov/22782721/ (Accessed March 12, 2025).

Filtz TM, Vogel WK, Leid M. 2013. Regulation of transcription factor activity by interconnected, post-translational modifications. Trends Pharmacol Sci 35: 76. https://pmc.ncbi.nlm.nih.gov/articles/PMC3954851/ (Accessed March 30, 2025).

Gil E, Hong SJ, Wu D, Park DH, Delgado RN, Malatesta M, Ahanger SH, Lin K, Villeda S, Lim DA. 2024. Chromatin regulator Kdm6b is required for the establishment and maintenance of neural stem cells in mouse hippocampus. Elife 13. https://elifesciences.org/reviewed-preprints/97262 (Accessed March 12, 2025).

Godfrey L, Crump NT, O’Byrne S, Lau IJ, Rice S, Harman JR, Jackson T, Elliott N, Buck G, Connor C, et al. 2020. H3K79me2/3 controls enhancer–promoter interactions and activation of the pan-cancer stem cell marker PROM1/CD133 in MLL-AF4 leukemia cells. Leuk 2020 351 35: 90–106. https://www.nature.com/articles/s41375-020-0808-y (Accessed March 25, 2025).

Greig LC, Woodworth MB, Galazo MJ, Padmanabhan H, Macklis JD. 2013. Molecular logic of neocortical projection neuron specification, development and diversity. Nat Rev Neurosci 14: 10.1038/nrn3586. https://pmc.ncbi.nlm.nih.gov/articles/PMC3876965/ (Accessed March 28, 2025).

Guo J, Walss-Bass C, Ludueña RF. 2010. The β isotypes of tubulin in neuronal differentiation. Cytoskeleton 67: 431–441. https://onlinelibrary.wiley.com/doi/full/10.1002/cm.20455 (Accessed March 27, 2025).

Harasym E, McAndrew N, Gomez G. 2017. Sub-micromolar concentrations of retinoic acid induce morphological and functional neuronal phenotypes in SK-N-SH neuroblastoma cells. Vitr Cell Dev Biol - Anim 53: 798–809. https://link.springer.com/article/10.1007/s11626-017-0190-x (Accessed March 28, 2025).

Hsieh J. Orchestrating transcriptional control of adult neurogenesis. http://www.genesdev.org/cgi/doi/10.1101/gad.187336.112. (Accessed March 28, 2025).

Huang Y, Zhang H, Wang L, Tang C, Qin X, Wu X, Pan M, Tang Y, Yang Z, Babarinde IA, et al. 2020. JMJD3 acts in tandem with KLF4 to facilitate reprogramming to pluripotency. Nat Commun 2020 111 11: 1–16. https://www.nature.com/articles/s41467-020-18900-z (Accessed March 12, 2025).

Huth JR, Bewley CA, Nissen MS, Evans JNS, Reeves R, Gronenborn AM, Clore GM. 1997. The solution structure of an HMG-I(Y)-DNA complex defines a new architectural minor groove binding motif. Nat Struct Biol 4: 657–665. https://pubmed.ncbi.nlm.nih.gov/9253416/ (Accessed March 12, 2025).

Imamura T, Izumi H, Nagatani G, Ise T, Nomoto M, Iwamoto Y, Kohno K. 2001. Interaction with p53 Enhances Binding of Cisplatin-modified DNA by High Mobility Group 1 Protein. J Biol Chem 276: 7534–7540.

Jacobson RH, Ladurner AG, King DS, Tjian R. 2000. Structure and function of a human TAFII250 double bromodomain module. Science 288: 1422–1425. https://pubmed.ncbi.nlm.nih.gov/10827952/ (Accessed March 25, 2025).

Jayaraman L, Moorthy NC, Murthy KGK, Manley JL, Bustin M, Prives C. 1998. High mobility group protein-1 (HMG-1) is a unique activator of p53. Genes Dev 12: 462. https://pmc.ncbi.nlm.nih.gov/articles/PMC316524/ (Accessed March 25, 2025).

Jessell TM. 2000. Neuronal specification in the spinal cord: inductive signals and transcriptional codes. Nat Rev Genet 1: 20–29. https://pubmed.ncbi.nlm.nih.gov/11262869/ (Accessed March 30, 2025).

Kar S, Patra SK. 2018. Overexpression of OCT4 induced by modulation of histone marks plays crucial role in breast cancer progression. Gene 643: 35–45. https://www.sciencedirect.com/science/article/pii/S0378111917310430?via%3Dihub#s0085 (Accessed September 12, 2025).

Kar S, Sengupta D, Deb M, Shilpi A, Parbin S, Rath SK, Pradhan N, Rakshit M, Patra SK. 2014. Expression profiling of DNA methylation-mediated epigenetic gene-silencing factors in breast cancer. Clin Epigenetics 6: 1–13. https://clinicalepigeneticsjournal.biomedcentral.com/articles/10.1186/1868-7083-6-20 (Accessed August 16, 2025).

Kirtana R, Manna S, Patra SK. 2023. KDM5A noncanonically binds antagonists MLL1/2 to mediate gene regulation and promotes epithelial to mesenchymal transition. Biochim Biophys Acta - Gene Regul Mech 1866: 194986.

Kovalevich J, Santerre M, Langford D. 2021. Considerations for the Use of SH-SY5Y Neuroblastoma Cells in Neurobiology. Methods Mol Biol 2311: 9–23.

Lange C, Huttner WB, Calegari F. 2009. Cdk4/cyclinD1 overexpression in neural stem cells shortens G1, delays neurogenesis, and promotes the generation and expansion of basal progenitors. Cell Stem Cell 5: 320–331. https://pubmed.ncbi.nlm.nih.gov/19733543/ (Accessed March 28, 2025).

Li W, Tian E, Chen ZX, Sun GQ, Ye P, Yang S, Lu D, Xie J, Ho TV, Tsark WM, et al. 2012. Identification of Oct4-activating compounds that enhance reprogramming efficiency. Proc Natl Acad Sci U S A 109: 20853–20858. https://pubmed.ncbi.nlm.nih.gov/23213213/ (Accessed March 27, 2025).

Liu Y, Xiao A. 2011. Epigenetic regulation in neural crest development. Birth Defects Res Part A Clin Mol Teratol 91: 788–796. https://onlinelibrary.wiley.com/doi/full/10.1002/bdra.20797 (Accessed March 28, 2025).

Liu Z, Chen O, Zheng M, Wang L, Zhou Y, Yin C, Liu J, Qian L. 2016. Re-patterning of H3K27me3, H3K4me3 and DNA methylation during fibroblast conversion into induced cardiomyocytes. Stem Cell Res 16: 507–518. https://pubmed.ncbi.nlm.nih.gov/26957038/ (Accessed March 12, 2025).

Maden M. 2007. Retinoic acid in the development, regeneration and maintenance of the nervous system. Nat Rev Neurosci 8: 755–765. https://pubmed.ncbi.nlm.nih.gov/17882253/ (Accessed March 30, 2025).

Mallaney C, Ostrander EL, Celik H, Kramer AC, Martens A, Kothari A, Koh WK, Haussler E, Iwamori N, Gontarz P, et al. 2019. Kdm6b regulates context-dependent hematopoietic stem cell self-renewal and leukemogenesis. Leukemia 33: 2506–2521. https://pubmed.ncbi.nlm.nih.gov/30936419/ (Accessed March 12, 2025).

Manna S, Kirtana R, Roy A, Baral T, Patra SK. 2023a. Mechanisms of hedgehog, calcium and retinoic acid signalling pathway inhibitors: Plausible modes of action along the MLL–EZH2-p53 axis in cellular growth control. Arch Biochem Biophys 742: 109600.

Manna S, Mishra J, Baral T, Kirtana R, Nandi P, Roy A, Chakraborty S, Niharika N, Patra SK. 2023b. Epigenetic signaling and crosstalk in regulation of gene expression and disease progression. Epigenomics 15: 723–740. https://pubmed.ncbi.nlm.nih.gov/37661861/ (Accessed March 12, 2025).

Margueron R, Reinberg D. 2011. The Polycomb complex PRC2 and its mark in life. Nat 2011 4697330 469: 343–349. https://www.nature.com/articles/nature09784 (Accessed March 12, 2025).

Matsui S, Granitto M, Buckley M, Ludwig K, Koigi S, Shiley J, Zacharias WJ, Mayhew CN, Lim HW, Iwafuchi M. 2024. Pioneer and PRDM transcription factors coordinate bivalent epigenetic states to safeguard cell fate. Mol Cell 84: 476–489.e10. https://www.cell.com/action/showFullText?pii=S1097276523010262 (Accessed March 27, 2025).

Miller SA, Mohn SE, Weinmann AS. 2010. Jmjd3 and UTX play a demethylase-independent role in chromatin remodeling to regulate T-box family member-dependent gene expression. Mol Cell 40: 594–605. https://pubmed.ncbi.nlm.nih.gov/21095589/ (Accessed April 1, 2025).

Mirizio G, Sampson S, Iwafuchi M. 2025. Interplay between pioneer transcription factors and epigenetic modifiers in cell reprogramming. Regen Ther 28: 246–252.

Navarro AI, Rico B. 2014. Focal adhesion kinase function in neuronal development. Curr Opin Neurobiol 27: 89–95. https://www.sciencedirect.com/science/article/pii/S095943881400052X (Accessed July 10, 2025).

Park DH, Hong SJ, Salinas RD, Liu SJ, Sun SW, Sgualdino J, Testa G, Matzuk MM, Iwamori N, Lim DA. 2014. Activation of Neuronal Gene Expression by the JMJD3 Demethylase Is Required for Postnatal and Adult Brain Neurogenesis. Cell Rep 8: 1290–1299.

Ramírez-González A, Ávila-López P, Bahena-Román M, Contreras-Ochoa CO, Lagunas-Martínez A, Langley E, Manzo-Merino J, Madrid-Marina V, Torres-Poveda K. 2023. Critical Role of the Transcription Factor AKNA in T-Cell Activation: An Integrative Bioinformatics Approach. Int J Mol Sci 24: 4212. https://pmc.ncbi.nlm.nih.gov/articles/PMC9965657/ (Accessed March 12, 2025).

Reeves R, Nissen MS. 1990. The A.T-DNA-binding domain of mammalian high mobility group I chromosomal proteins. A novel peptide motif for recognizing DNA structure. J Biol Chem 265: 8573–8582.

Sanchez AM, Shortrede JE, Vargas-Roig LM, Flamini MI. 2016. Retinoic acid induces nuclear FAK translocation and reduces breast cancer cell adhesion through Moesin, FAK, and Paxillin. Mol Cell Endocrinol 430: 1–11. https://www.sciencedirect.com/science/article/pii/S0303720716301344?via%3Dihub#sec3 (Accessed July 10, 2025).

Sansom SN, Griffiths DS, Faedo A, Kleinjan DJ, Ruan Y, Smith J, Van Heyningen V, Rubenstein JL, Livesey FJ. 2009. The level of the transcription factor Pax6 is essential for controlling the balance between neural stem cell self-renewal and neurogenesis. PLoS Genet 5. https://pubmed.ncbi.nlm.nih.gov/19521500/ (Accessed March 27, 2025).

Schaefer T, Mittal N, Wang H, Ataman M, Candido S, Lötscher J, Velychko S, Tintignac L, Bock T, Börsch A, et al. 2024. Nuclear and cytosolic fractions of SOX2 synergize as transcriptional and translational co-regulators of cell fate. Cell Rep 43. https://pubmed.ncbi.nlm.nih.gov/39368083/ (Accessed March 28, 2025).

Sengupta D, Deb M, Rath SK, Kar S, Parbin S, Pradhan N, Patra SK. 2016. DNA methylation and not H3K4 trimethylation dictates the expression status of miR-152 gene which inhibits migration of breast cancer cells via DNMT1/CDH1 loop. Exp Cell Res 346: 176–187. https://www.sciencedirect.com/science/article/pii/S0014482716302117?via%3Dihub#s0080 (Accessed September 12, 2025).

Shait Mohammed MR, Zamzami M, Choudhry H, Ahmed F, Ateeq B, Khan MI. 2022. The Histone H3K27me3 Demethylases KDM6A/B Resist Anoikis and Transcriptionally Regulate Stemness-Related Genes. 10: 780176. https://pmc.ncbi.nlm.nih.gov/articles/PMC8847600/ (Accessed March 12, 2025).

Siddiqa A, Sims-Mourtada JC, Guzman-Rojas L, Rangel R, Guret C, Madrid-Marina V, Sun Y, Martinez-Valdez H. 2001. Regulation of CD40 and CD40 ligand by the AT-hook transcription factor AKNA. Nature 410: 383–387. https://pubmed.ncbi.nlm.nih.gov/11268217/ (Accessed March 12, 2025).

Stolerman ES, Francisco E, Stallworth JL, Jones JR, Monaghan KG, Keller-Ramey J, Person R, Wentzensen IM, McWalter K, Keren B, et al. 2019. Genetic variants in the KDM6B gene are associated with neurodevelopmental delays and dysmorphic features. Am J Med Genet Part A 179: 1276–1286. https://onlinelibrary.wiley.com/doi/full/10.1002/ajmg.a.61173 (Accessed March 28, 2025).

Svotelis A, Bianco S, Madore J, Huppé G, Nordell-Markovits A, Mes-Masson AM, Gévry N. 2011. H3K27 demethylation by JMJD3 at a poised enhancer of anti-apoptotic gene BCL2 determines ERα ligand dependency. EMBO J 30: 3947–3961. https://pubmed.ncbi.nlm.nih.gov/21841772/ (Accessed April 1, 2025).

Swigut T, Wysocka J. 2007. H3K27 demethylases, at long last. Cell 131: 29–32. https://pubmed.ncbi.nlm.nih.gov/17923085/ (Accessed March 12, 2025).

Taberlay PC, Kelly TK, Liu CC, You JS, De Carvalho DD, Miranda TB, Zhou XJ, Liang G, Jones PA. 2011. Polycomb-Repressed Genes Have Permissive Enhancers that Initiate Reprogramming. Cell 147: 1283–1294.

Takahashi K, Yamanaka S. 2006. Induction of Pluripotent Stem Cells from Mouse Embryonic and Adult Fibroblast Cultures by Defined Factors. Cell 126: 663–676. https://pubmed.ncbi.nlm.nih.gov/16904174/ (Accessed March 27, 2025).

Tan BT, Wang L, Li S, Long ZY, Wu YM, Liu Y. 2015. Retinoic acid induced the differentiation of neural stem cells from embryonic spinal cord into functional neurons in vitro. Int J Clin Exp Pathol 8: 8129. https://pmc.ncbi.nlm.nih.gov/articles/PMC4555709/ (Accessed March 28, 2025).

Trapnell C, Cacchiarelli D, Grimsby J, Pokharel P, Li S, Morse M, Lennon NJ, Livak KJ, Mikkelsen TS, Rinn JL. 2014. The dynamics and regulators of cell fate decisions are revealed by pseudotemporal ordering of single cells. Nat Biotechnol 2014 324 32: 381–386. https://www.nature.com/articles/nbt.2859 (Accessed March 29, 2025).

Wang W, Cho H, Lee JW, Lee SK. 2022a. The histone demethylase Kdm6b regulates subtype diversification of mouse spinal motor neurons during development. Nat Commun 2022 131 13: 1–22. https://www.nature.com/articles/s41467-022-28636-7 (Accessed March 12, 2025).

Wang Y, Khandelwal N, Liu S, Zhou M, Bao L, Wang JE, Kumar A, Xing C, Gibson JR, Wang Y. 2022b. KDM6B cooperates with Tau and regulates synaptic plasticity and cognition via inducing VGLUT1/2. Mol Psychiatry 27: 5213–5226. https://pubmed.ncbi.nlm.nih.gov/36028572/ (Accessed March 12, 2025).

Wang Z, Oron E, Nelson B, Razis S, Ivanova N. 2012. Distinct lineage specification roles for NANOG, OCT4, and SOX2 in human embryonic stem cells. Cell Stem Cell 10: 440–454. https://pubmed.ncbi.nlm.nih.gov/22482508/ (Accessed March 28, 2025).

Wiederschain D, Kawai H, Gu J, Shilatifard A, Yuan Z-M. 2003. Molecular Basis of p53 Functional Inactivation by the Leukemic Protein MLL-ELL. Mol Cell Biol 23: 4230. https://pmc.ncbi.nlm.nih.gov/articles/PMC156137/ (Accessed March 25, 2025).

Williams K, Christensen J, Rappsilber J, Nielsen AL, Johansen JV, Helin K. 2014. The histone lysine demethylase JMJD3/KDM6B is recruited to p53 bound promoters and enhancer elements in a p53 dependent manner. PLoS One 9. https://pubmed.ncbi.nlm.nih.gov/24797517/ (Accessed March 12, 2025).

Xie HR, Hu L Sen, Li GY. 2010. SH-SY5Y human neuroblastoma cell line: In vitro cell model of dopaminergic neurons in Parkinson’s disease. Chin Med J (Engl) 123: 1086–1092.

Yang J, Milasta S, Hu D, AlTahan AM, Interiano RB, Zhou J, Davidson J, Low J, Lin W, Bao J, et al. 2017. Targeting Histone Demethylases in MYC-Driven Neuroblastomas with Ciclopirox. Cancer Res 77: 4626–4638. https://pubmed.ncbi.nlm.nih.gov/28684529/ (Accessed March 12, 2025).

Yeo JC, Ng HH. 2012. The transcriptional regulation of pluripotency. Cell Res 2013 231 23: 20–32. https://www.nature.com/articles/cr2012172 (Accessed March 27, 2025).

Young RA. 2011. Control of the embryonic stem cell state. Cell 144: 940–954. https://www.cell.com/action/showFullText?pii=S0092867411000717 (Accessed March 27, 2025).

Yu J, Vodyanik MA, Smuga-Otto K, Antosiewicz-Bourget J, Frane JL, Tian S, Nie J, Jonsdottir GA, Ruotti V, Stewart R, et al. 2007. Induced pluripotent stem cell lines derived from human somatic cells. 318: 1917–1920. https://pubmed.ncbi.nlm.nih.gov/18029452/ (Accessed March 27, 2025).

Yun J, Kim YS, Jung JH, Seo PJ, Park CM. 2012. The AT-hook Motif-containing Protein AHL22 Regulates Flowering Initiation by Modifying FLOWERING LOCUS T Chromatin in Arabidopsis. J Biol Chem 287: 15307. https://pmc.ncbi.nlm.nih.gov/articles/PMC3346147/ (Accessed March 25, 2025).

Zhao W, Li Q, Ayers S, Gu Y, Shi Z, Zhu Q, Chen Y, Wang HY, Wang RF. 2013. Jmjd3 inhibits reprogramming by upregulating expression of INK4a/Arf and targeting PHF20 for ubiquitination. Cell 152: 1037–1050. https://pubmed.ncbi.nlm.nih.gov/23452852/ (Accessed March 12, 2025).

